# Estimation of Task-Evoked Directed Functional Connectivity by Cross-Mapping Psychophysiological Variables

**DOI:** 10.1101/2022.10.21.513137

**Authors:** Ameer Ghouse, Johannes Schultz, Gaetano Valenza

## Abstract

Understanding the functional connectivity between different brain regions is vital for improving our comprehension of neural processing and cognition. While directed functional connectivity methods can provide us with statistical estimates of information exchange between regions, classic exploratory methods may not capture the nonlinear temporal effects that are observed in fMRI-BOLD data during task-evoked neural activity. To address this limitation, we propose a novel methodology that leverages variational cross-mapping analysis, inspired by psychophysiological interactions, to identify directional influence between connected regions of interest. Our approach can help uncover previously unknown patterns of information exchange and account for nonlinear effects, making it a valuable addition to the toolkit of researchers studying brain function. We demonstrate the effectiveness of our method using simulated neurovascular signals and publicly available fMRI data from 680 human participants performing an emotional face processing task. Our results suggest information flows from the occipital face area to the superior temporal sulcus and the fusiform face area, and additionally from the superior temporal sulcus to the fusiform gyrus. These findings are consistent with previously documented effective connectivity findings in face processing and provide new insights into the exploratory analyses of non-linear directed connectivity for task-evoked data. Overall, our findings contribute to advancing our understanding of directed functional connectivity in the brain and demonstrate the potential of our method to uncover previously unknown patterns of information exchange.

**Author summary:** The advent of large datasets has made it possible for many research groups to explore functional connectivity between different brain regions. The ability to assess directed connectivity between multiple regions from task-evoked neural responses could potentially uncover connections that were not previously hypothesized based on available data. However, classic methods for exploring task-evoked effects often rely on specific assumptions that are frequently violated by the data, such as nonlinearity, stationarity, and separability of cause from effect.

Recent studies have attempted to address these issues using sliding window approaches or parameterized forward causal models, but these methods have limitations such as fixed contextual effect windows or restricted search space for forward models. To overcome these challenges, we propose a Bayesian non-parametric cross-mapping method that can address non-linearity and separability while using specially designed covariance functions to address non-stationarity.

We demonstrate through simulations that our proposed method can detect pair-wise interacting neural populations with high sensitivity and specificity, and accurately infer changes in connections between tasks in both acyclical and cyclical neural networks. We also show that our method can replicate known connectivity findings about emotional face processing in a publicly available dataset. Thus, our method represents a promising exploratory connectivity tool for cognitive and behavioral neurosciences.

## 1 Introduction

Exploring task-evoked coupling between neural structures can be fraught with complications, particularly when coupling occurs in latent variables as is the case with functional magnetic resonance imaging (fMRI) blood oxygen level dependent (BOLD) data. How can we determine coupling effects evoked by a particular stimulus in a system, when residual effects of prior task-related events are overlapping? This is an especially important question in functional coupling analyses involving contrasting of task-evoked signals. Such analyses are often performed in human neuroimaging to understand how coupling between activity in different structures changes between a phenomenon of interest (often, a psychologically-defined experimental condition) and a control activity. Further complications arise due to functional coupling in neuroimaging data being not only nonlinear but complex due to the high dimensionality of the data, a consequence of the many interactions and feedback loops that constitute neural circuits [1–4].

Coupling analysis in neuroimaging, hereon referred to as connectivity, can be divided into two camps: functional and effective connectivity. Functional connectivity hinges on discovering statistical relationships between regions of interest, such as correlation [5, 6]. This kind of connectivity analysis, however, is prone to confounds, such as violations of Gaussianity, which may lead to false positives [7, 8]. Partial correlation may help by performing conditional correlation tests, either by regressing out a confounder using a general linear model (GLM) – i.e. a linear regression that assumes independent, normally distributed residuals – or through the precision matrix (inverse of the covariance matrix) of the voxels in a functional image [9]. However the conditional tests that quantify such partial effects embed no information of temporality through which physical interactions operate, and may thus underestimate effects or in some cases yield false positive results, due to non-independence of the data acquired at successive time-points.

Some methods assess functional connectivity in a directed manner. Psychophysiological interaction (PPI) is one of these methods [10–12]. It exploits the General Linear Model (GLM) and so-called psychophysiological interaction variables: time series constructed from deconvolved BOLD data that are element-wise multiplied with a vector describing psychological events that occur in an experimental run. PPI aims to uncover brain regions whose functional connectivity with a seed region changes depending on the occurrence of a psychological variable of interest. However, this method takes no consideration of the direction of connectivity.

On the other hand, effective connectivity attempts to ascertain the mechanistic properties of a network, or in other words the influence of one neural system on another [6]. Dynamic Causal Modeling (DCM) is a specific effective connectivity method that can take into consideration the nonlinear interactions in time of a system, and predict how coupling may emerge from it. However, this method involves an exhaustive combinatorial search through candidate models in order to select the one with the most evidence given a-priori expectations [13]. There have been developments of the DCM framework to make it more appropriate for whole-brain exploratory analysis, particularly regression DCM [14]. However, this method sacrifices nonlinear characterizations of coupling in favour of computational feasibility.

An intermediate approach between effective connectivity methods such as DCM and functional connectivity methods like partial correlation is directed functional connectivity. Granger Causality (GC), and its nonparametric counterpart transfer entropy, are members of this approach. These methods take temporal precedent information into consideration. However, these methods are inherently memory-less, and can thus not handle nonlinear and temporal context-dependent effects observed in neural data [13, 15–18]. Although directed functional connectivity is more appropriate for performing exploratory analysis, these methods hinge on stationary assumptions, and thus require the use of a sliding window approach to infer task-evoked activity. Therefore, such methods may ignore crucial effects that emerge from dynamic complex effects such as memory, as determined by location in state space. Furthermore, GC and transfer entropy can be prone to reveal false positive directions of influence [19, 20].

In this paper, we aim to detail the theory and implementation of a method that can infer directed functional connectivity while taking into account the non-stationary properties of a task-evoked response, eschewing the sliding window approach. In brief, the proposed method exploits the ideas of state space reconstruction embedding methods [21, 22], Gaussian Processes [23], and psychophysiological interactions [10–12] to develop a generative (probabilistic) model of the task-evoked response coupling activity. From this probabilistic model, we can then perform Bayesian inference to determine which direction of coupling provides more evidence for the observed data. The proposed model’s theory has its foundations in the idea of cross-mapping [24], and has already shown promise as a robust measure of connectivity in the non-task evoked case in our prior papers introducing Gaussian Process Convergent Cross Mapping (GP-CCM) [25, 26]. We will further show in this paper that the Bayesian modeling provides further flexibility to account for mediating factors in path analysis. An added benefit of the method is that it is entirely non-parametric, i.e. the model is a function of the data.

We test the face validity of the method using simulations of hemodynamic responses evoked by interacting brain regions in pairwise configurations, as well as in acyclic and cyclic network topologies. The construct validity of the method is then verified using a publicly available face processing BOLD dataset [28]. We hypothesized that from this dataset, our proposed method would be able to replicate network interactions findings in the extensively studied face processing system [29, 30, 32–36]. Core regions of the face processing system include face-sensitive parts of the inferior occipital gyrus (such as the Occipital Face Area or OFA), the fusiform gyrus (FFA) and of the superior temporal sulcus (STS). Overall, previous findings suggest that these three core regions form two parallel connection streams: a ventral stream connecting OFA to FFA (and onwards to other regions particularly sensitive to static information, such as face identity), and a dorsal stream connecting OFA to STS (and onwards to other regions sensitive to dynamic or variable information, such as facial expression). Connections are thought to run both forward and backward within each stream. While this simplistic description summarises many structural and functional connectivity findings, some studies report a high degree of functional interconnection between all three regions, particularly in the right hemisphere (e.g., [36])). The amount of documented findings about this network makes it an ideal test bed to evaluate our proposed method.

## 2 Materials and methods

This section is organised as follows: First, we will briefly introduce the theory of cross-mapping methods. Next, we will describe how we extend the method to task-evoked interactions and how to apply it to neurovascular signals. Then we describe our face-validity tests for uncovering network interactions. Finally, we describe the construct validation tests that we performed on a publicly available human BOLD dataset.

### 2.1 Background on Embeddings and the Cross-Mapping Methods (CCM)

Embedding theorems such as Takens’ allow reconstruction of state space variables in a higher dimensional space [21, 22], for example using lag coordinates:

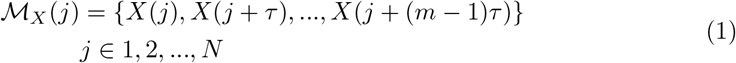

Where X is the time series (which we assume is generated from some dynamical system 𝒳), j is a time index, ℳ_*X*_ represents the reconstructed phase space of 𝒳 from X, *m* and *τ* respectively are the embedding dimension and the delay-time for the state-space reconstruction, and *N* + (*m*− 1)*τ* are the amount of time points in the time series. Accordingly, we can encode the embedding vectors into a ℝ^*N ×m*^ matrix [25] as follows:

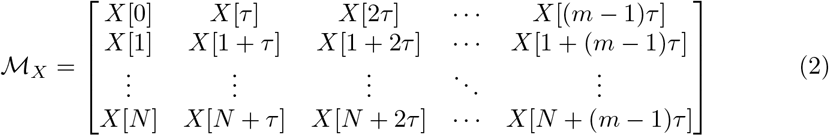

We use ℳ_*X*_ (*j, s*) to specify the element at the *j*^*th*^ row and the *s*^*th*^ column, and ℳ_*X*_ (*j*) to refer to the vector resulting from the *j*^*th*^ row.

Originally, convergent cross-mapping (CCM) in Sugihara et al [24] leveraged such reconstructed phase spaces to devise a measure of coupling for dynamical systems based on cross-prediction. While CCM may measure directed coupling between two dynamical systems [24], this model does not incorporate measurement or process noise. To overcome this limitation, we previously expanded upon the idea of cross-mapping by considering the problem as determining information gain using Bayesian inference, introducing Gaussian Process Convergent Cross-Mapping (GP-CCM) [25].

A Gaussian process can be defined by a mean and covariance function [23], i.e. 𝒢 𝒫 (*m*(*x*), *K*(*x, x*^*′*^)). Considering our interest in primarily second-order statistics and the fact that we can subtract the mean signal from data, we define *m*(*x*) as a zero mean function. For the covariance, we specify a squared exponential radial basis function between points on the phase space as follows:

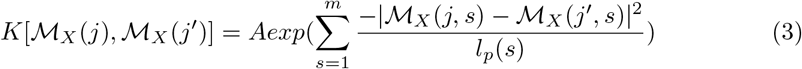

Where *A* encodes the magnitude of the marginal variances, and *l*_*p*_(*s*) encodes how sensitive the distance function is for the column *s* of the embedding vector [37].

#### 2.1.1 Incorporating Psychophysiological Interactions into Gaussian Process CCM

In this study, we introduce how to encode specificity for task-evoked interactions in the covariance function shown in eq. 3 by using psychophysiological interaction (PPI) variables [10].

Psychophysiological interactions originated as a model that exploits the design matrix of a general linear model (GLM) to infer task-evoked coupling between regions of interest. However, there is no embedding of dynamical or nonlinear information in PPI. We propose our method, Psychophysiological Interactions based Gaussian Process Convergent Cross-Mapping (PPI GP-CCM), as an extension of PPI methods to overcome this limitation.

To begin, consider interactions between voxels (or surface vertices). We denote the *i*^*th*^ Psychophysiological Interactions (*PP*_*i*_) between the *k*^*th*^ volumetric BOLD voxel (or surface vertex) *B*_*k*_ and *i*^*th*^ class of events as the following element-wise product:

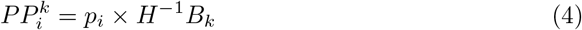

where *H* is the hemodynamic response function in a Toeplitz matrix form [12], and the *p*_*i*_ vector designates the *i*^*th*^ class of events by setting values to 1 at the time points of psychological events of interest, and -1 everywhere else [38].

Here, we exploit the aforementioned formalization of the *i*^*th*^ psychophysiological interaction (the *i*^*th*^ task-evoked interaction of interest) with the *k*^*th*^ volumetric BOLD voxel, i.e. 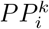, and incorporate it into the a-priori covariance function in 3, i.e., 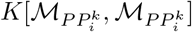. We account for measurement noise as i.i.d. Gaussian random variables with zero mean and standard deviation *σ*. To further embed the *i*^*th*^ psychophysiological interaction 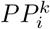 into a nonlinear kernel 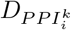, we define the a-priori covariance matrix of the Gaussian process underlying each voxel or surface vertex *k* as follows:

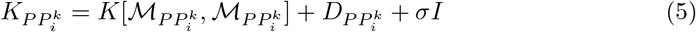

where *D* is a *NxN* diagonal matrix corresponding to the *N* samples of the reconstructed phase space. Of note, *D* biases the diagonal elements of 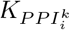 to associate larger marginal variance when a participant is not engaging in the *i*^*th*^ task (see Fig. 1 for an illustration). In particular, the diagonal bias *d* term at time point *j* for the covariance as:

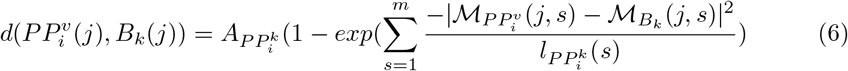

As the distance between the psychophysiological interaction variable and the original BOLD signal decreases, such a diagonal component tends towards zero. As it increases, it tends towards towards a saturating value 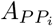. Eq. 6 lays out the foundation to take into consideration nonstationary mediating factors of the temporal BOLD signal by performing the calculation for each time sample, i.e., for each *j*. For an illustration of this task-biasing factor for the covariance function, see Fig. 1.

**Fig 1.**
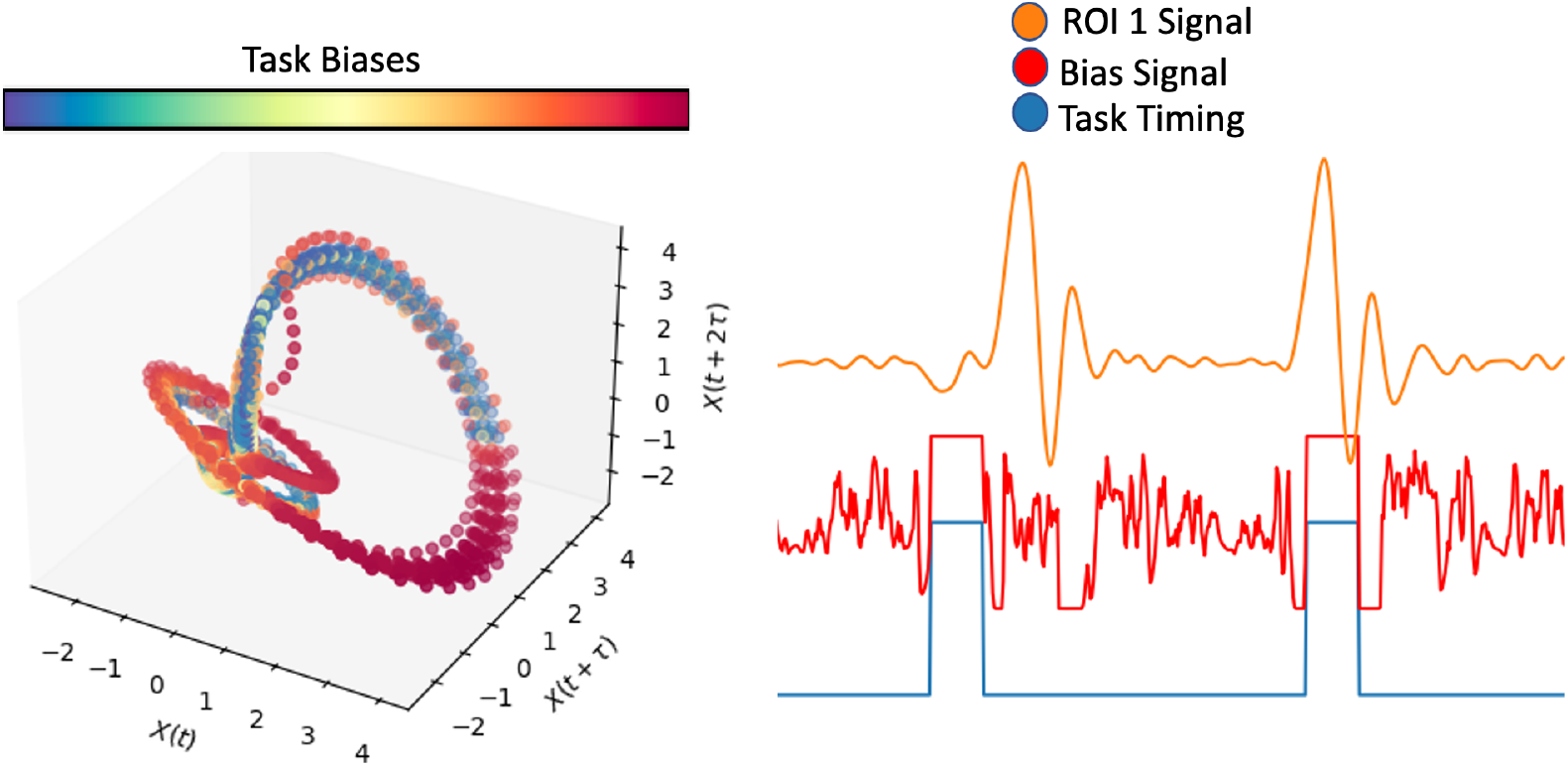
On the left, an embedding of a BOLD signal evoked from eq. 11 is shown, where the states on the left are colorcoded according to their “task-biases”, resulting from the task-similarity function in eq. 6, where warmer colors represent higher similarity, and colder colors represent lower similarity. On the right, time series are shown with the original BOLD signal in orange, the task-biasing signal for the diagonal of the cross-mapping covariance in red, and the task-timing signal in blue.

In many analyses of neuroimaging data, we are interested in the path of interactions that yield a functional response. This is the precise reason why partial correlation measures are used in functional connectivity [9] and why Granger Causality exploits multivariate autoregressive models [39–41]. It is also why DCM takes into consideration the full connectivity matrix in its bilinear state equations for neural metabolic demand [42, 43]. Similarly, for the proposed method, we must take into consideration the mediating factors inside the model. Thus, we can further extend eq. 5 for all possible mediating BOLD signals during posterior inference. The a-priori covariance matrix of the Gaussian process model testing for the cross-mapping effect from voxel or surface vertex *r* onto *k* would follow as:

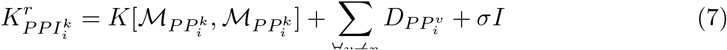

As in [25], using the methods described in [44], sparsity is then enforced on this a priori covariance through a Nystrom approximation to reduce the dimensionality of the a-priori covariance by exploiting representative pseudopoints *Z*. We can obtain the task-evoked posterior covariance of cross-mapping voxel *r* onto voxel *k* for event *i*, i.e. 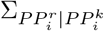 by exploiting the definition of the conditional multivariate Gaussian distributions [23]:

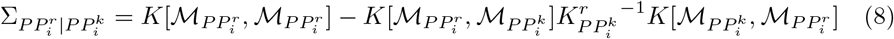

The posterior covariance 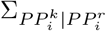 for cross-mapping voxel *k* onto voxel *r* can be obtained similarly. The coupling strength can then be determined by exploiting entropies provided by the posterior covariances.

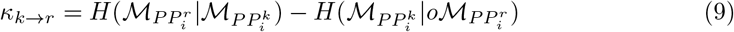

Negative values of *κ*_*k*→*r*_ indicate directed connectivity in the opposite direction. The entropy terms are derived from the definition of the differential entropy of Gaussian distributions [26, 27]:

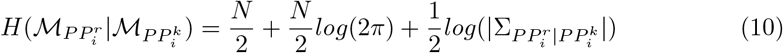

A variational Bayesian method outlined in [26] is used to integrate the effects of hyperparameters out of the connection strength statistic through approximations of posterior distributions of the hyperparameters via Gaussian distributions.

The proposed method can further be used in region of interest (ROI) analysis by reconstructing a state space using concatenations of delay coordinates constructed for individual time series. If *ROI*_1*v*_ specifies the *v*^*th*^ voxel of *ROI*_1_, then 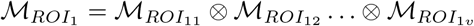, where ⊗ denotes a concatenation. The dimensionality of state spaces scales linearly with the amount of voxels in the ROI, thus these state spaces can quickly reach high dimensions, rendering radial basis functions less sensitive [45]. Dimensionality reduction techniques such as principal component analyses (PCA) are then recommended to reduce the dimensionality of these state spaces.

The hypothesis test for PPI GP-CCM is *H*_*o*_ : 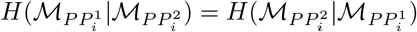 with 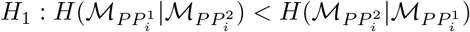 if *ROI*_1_ directs information towards *ROI*_2_ during event i. The null distribution is constructed using a permutation analysis as described in [25] and [26]. We use *α* to designate the significance level for rejecting the null hypothesis at the individual subject level.

### 2.2 Face Validity Tests

Face validity tests correspond to validation measures that ensure that the proposed method can achieve its stated goals in controlled environments [46]. Here, face validity comprises simulations, particularly for BOLD signals evoked by latent interactions in neural networks in acyclic topologies described by bilinear neural state equations [42]. These simulations test whether the proposed method can identify characteristic differences between conditions. In the supplementary material, we also provide simulations of cyclic topology configurations, which are common in neural data obtained during cognitive processes [47–50], thus important to face-validate. also includes receiver operating characteristic curve analyses to assess sensitivity and specificity of the method, as well as an illustration of how the proposed statistic successfully follows connectivity strength trends.

Furthermore, in the supplementary material, we further show results on simulations of pairwise-coupled populations of neurovascular neural networks. These simulations aimed to test how different inter-event time intervals affect the measures obtained by our proposed method. We furthermore varied ground truth connectivity to assess whether the trends of inferred connection strengths follow the ground truth. Finally, for sensitivity and specificity analysis, we performed receiver operating characteristic curve analyses.

In order to assess the validity of the proposed PPI GP-CCM on metabolic signals related to neural activity as obtained using functional Near Infrared Spectroscopy (fNIRS) or functional Magnetic Resonance Imaging (fMRI), we exploit forward models of hemodynamics to simulate neurovascular signals emerging from neural networks [43]. These models reflect the nonlinear behavior of hemodynamics evoked by neural activity. In these simulations, we have 3 populations, each comprising 50 neural units for a total of 150 neural units. As per [43], a bilinear state equation is used to model neural activity signals (*x*) induced by arbitrary psychological events:

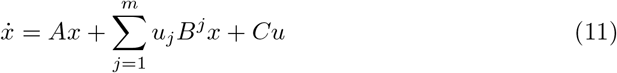

The *A* matrix describes the autonomous dynamics of the system; in our simulations, we treat it as a diagonal matrix with values -1, implying dynamics returning to a baseline 0 state. *u* are psychological events (where the element *u*_*j*_ is 0 at time points when *jth* condition is not occurring, and 1 at time points when the *j*^*th*^ condition is occurring). In our experiments, we have 2 conditions, plus a baseline condition of no cross-population interactions, thus *j*∈ 1, 2, 3.

*B*^*j*^ describes the interconnections between populations of interest. For the acyclic topology shown in Fig. 3, the B matrix would be 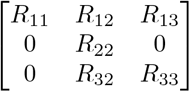. Each *R*_*ij*_ is an i.i.d. random matrix *R*_*ij*_ ∼ ℳ *𝒩* (0, 0.4, 1). The B matrices do not change in time, rather they are realized for each simulated time course. *C* describes which populations the stimulus condition interacts with, which in this paper is simulated as a diagonal matrix. *u*_3_ in all simulations is a binomial point process with *p* = 0.3 that describes random neural activity that elicit no functional interconnections between populations. This serves as background neural noise on top of the activities.

**Fig 2.**
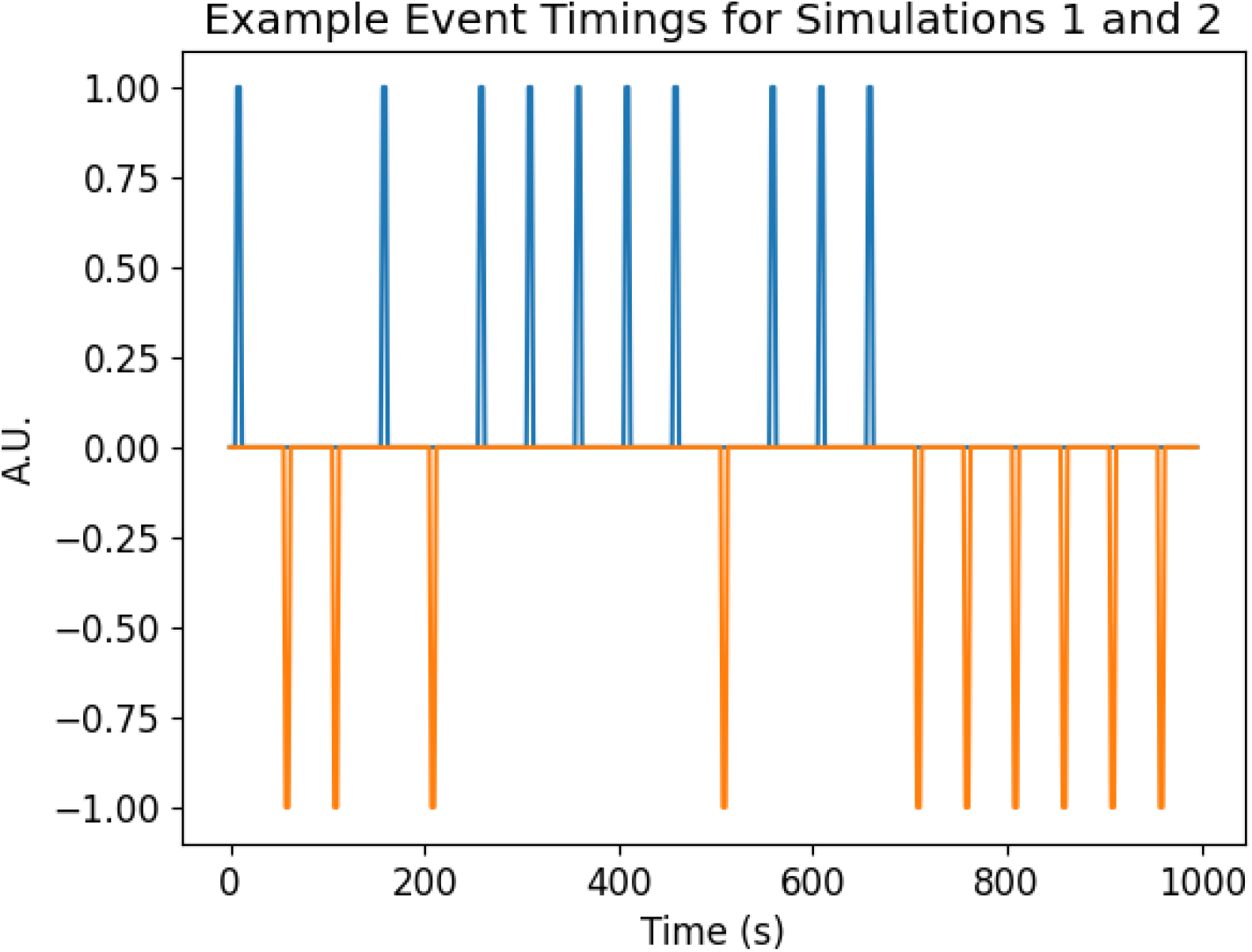
Event timings for the conditions being simulated

**Fig 3.**
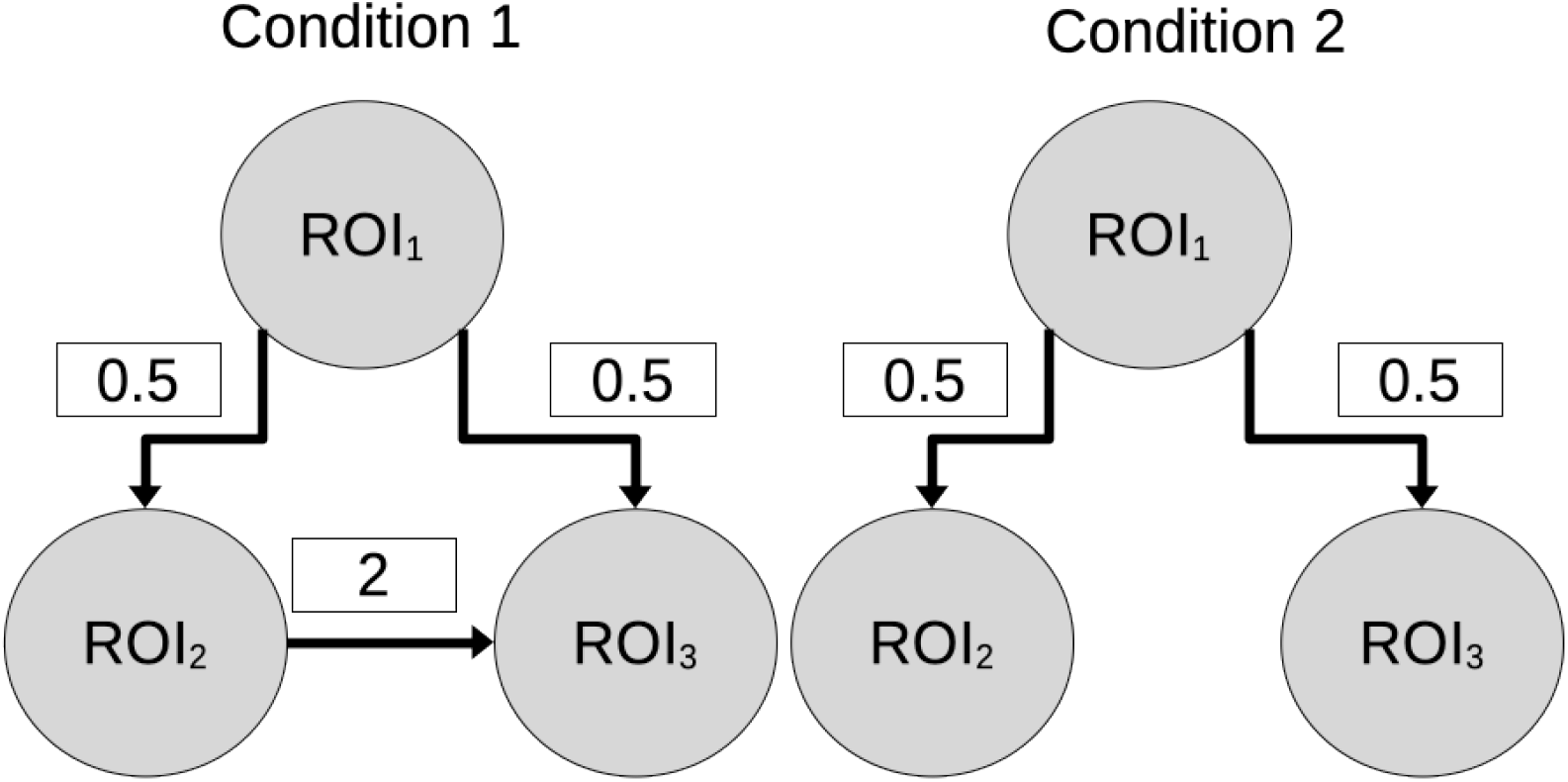
We simulate network with acyclic functional interactions. In condition 1, the connection between *ROI*_2_ and *ROI*_3_ is closed (information can flow), while in condition 2, the connection is open (information can not flow).

After the neural state equations, regional blood flow *f*, blood volume *v*, and deoxyhemoglobin concentration *q* is simulated using the parameters and mechanistic equations proposed in [42, 43], also seen in table 1. In other words, parameters of the hemodynamic model were independently sampled from Gaussian distributions for each simulated voxel according to table 1, following parameter values specified in [42]. A noise corrupted deoxyhemoglobin time series vector *q*^*^ = *q* + *ϵ*, where *ϵ* is additive noise sampled from 𝒩 (0, 0.1*I*), is saved for further processing. This additive noise process should give the simulations a mean signal-to-noise ratio (SNR) of 4 for the neural population. Processing after the simulations comprises deconvolution, PCA dimensionality reduction of networks, and PPI GP-CCM analysis.

**Table 1.** Prior distributions of hemodynamic generating parameters for the simulations, courtesy of [42]

| Parameter | Description | Prior Mean $\mu$ | Prior Variance $\sigma$ |
| --- | --- | --- | --- |
| $\kappa$ | Rate of signal decay | 0.65 per s | 0.015 |
| $\tau_f$ | Rate of flow-dependent elimination | 0.41 per s | 0.002 |
| $\tau_v$ | Hemodynamic transit time | 0.98 s | 0.0568 |
| $\alpha$ | Grubb's exponent | 0.32 | 0.0015 |
| $E_o$ | Resting oxygen extraction fraction | 0.34 | 0.0024 |

#### 2.2.1 Simulating Network Topologies of Neural Populations

In real brains, more than two neural populations interact at a given time. Thus, great care needs to be taken to consider mediating factors when making inferences about connectivity. To test our model on network interactions with more than two populations, we consider two kinds of topologies (see Fig. 3 and Supplementary Fig. 4). The first topology (Fig. 3) is an acyclic graph, where directed interactions follow the assumptions of the PPI GP-CCM model: unidirectional interactions exist between all nodes with no recurrence. In the supplementary material, we also simulate cyclic topologies (Supplementary Fig. 4 and 5) where recurrences can occur. We simulate BOLD time series for the networks with 2 conditions: experimental condition 1 where all connections in the network are closed (information flows through all connections), and condition 2 where the connection described between *ROI*_2_ and *ROI*_3_ is severed or opened (information can not flow). We exploit eq. 7 to control for mediated interactions to uncover the network structure. We hypothesize that cutting the connection between two nodes (ROIs linked by the dashed arrow in Fig. 3) would result in a significantly smaller measured connectivity between them.

Applying an analysis pipeline similar to those used in empirical datasets [56], we want to contrast the functional network topology observed in conditions 1 and 2. In order to do so, we perform a paired Wilcoxon signed rank test between the inferred connection strengths in the two conditions to determine which edge in the network graph shows significant differences between the two conditions.

### 2.3 Construct Validity Tests

Construct validity tests are analyses used to confirm that the proposed method will achieve its stated goals when faced with an environment where it would be expected to be used in [57]. In this paper, we exploit a large dataset from the WU-Minn HCP Consortium S900 Release [28] where individuals are asked to differentiate between emotional faces and arbitrary shapes [58]. Given the established literature regarding feedforward face perception circuitry, we focus on contrasting connectivities evoked when presenting face vs. shape stimuli to replicate previously published effects [29–31, 35]. The data are publicly available at https://www.humanconnectome.org.

#### 2.3.1 Validation of PPI GP-CCM Using a Face Perception Dataset

We perform PPI GP-CCM to assess the connectivity between data from ROIs specified to be face-sensitive according to the face functional localizer study performed in [35] for the human connectome project S900 dataset. The dataset was preprocessed using NiLearn’s clean img function, which comprises detrending using cosine basis functions, high-pass filtering (0.008 Hz cutoff frequency), and movement confound removal. Afterwards, the functional images were smoothed by a 4mm FWHM Gaussian filter. Consistent with [35], we only use subjects in the HCP dataset with a robust signal in a separate task (working memory, n-back task [59]) for decoding between face stimuli vs other stimuli in the face localized ROIs, thus resulting in a sample size of N=680.

We aimed to use our proposed connectivity method to qualitatively replicate effective connectivity findings in important regions of the network underlying face processing: the early visual cortex (EVC), the occipital face-processing region (OFA), face-sensitive part of the fusiform gyrus (FFA) and the face-sensitive part of the cortex in the superior temporal sulcus (STS). Structural connections between OFA and FFA are well known [32]. DCM-based effective connectivity analyses of fMRI data [29] and of combined fMRI-EEG data [60] suggest that these regions are organized as a feed-forward architecture in which the EVC and OFA have a direct influence on the FFA and STS, while the STS has direct influence on the FFA. We thus expected that faces would evoke greater directed connectivity strength from occipital areas to the FFA and STS compared to every other type of stimulus in the dataset. Functional connectivity from the superior temporal areas to the fusiform areas would not change as much as OFA-FFA and OFA-STS.

To apply the PPI GP-CCM analysis to our data, we first deconvolved the BOLD signal mapped to the surface vertices using a Fourier set and the hemodynamic response function described by Glover et al. [51]. The Fourier coefficients are determined in a regularized maximum a-posteriori (MAP) approach defined in [12]. Concretely, we determine the coefficients for a linear time-invariant system:

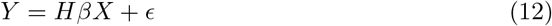

where H is the hemodynamic response convolution matrix in Toeplitz form, *β* are the Fourier coefficients, X is the matrix of Fourier bases, and *ϵ* is the error between the observed BOLD signal Y and the deconvolution model. The MAP solution to this system is:

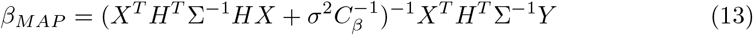

where *C*_*β*_ is a prior precision matrix, controlling the frequencies to attenuate in the deconvolution solution, ∑ is the error correlation matrix, and *σ* is the error variance. Ordering the rows of the bases by ascending frequency (f), we define the a-priori precision matrix with diagonal components equal to 1*/f* in order to prefer solutions concentrated in lower frequencies.

From the deconvolved BOLD signal, we make 2 psychophysiological interaction variables, similar to the approach used in the gPPI toolbox [61], based on the timestamps of the face and shape events. We use the cluster centers for the face localized ROIs specified in Supplementary table 2 in [35] as the representative BOLD signal time course for each ROI for performing PPI GP-CCM analysis. To assess whether the connections are significantly stronger when viewing faces compared to houses in the surviving connections for face perception, we use a one-sided paired T tests on the group level (N=680), calculated for each ROI to ROI interaction to determine which interactions had significantly greater effect when a participant was presented with a face stimuli. The results of these tests were considered significant if the obtained p value was *<* 0.05 after correcting for 16 multiple tests using a Bonferroni correction.

## 3 Results

### 3.1 Face Validity Tests

For face validity, we assessed connectivity in simulated topological network interactions of neural populations as described on section 2.2.1 and illustrated in Fig. 3. We assessed whether contrast statistics, i.e. the Wilcoxon signed rank test, are able to detect changes in connections strengths between two conditions, over the 100 simulations. For cyclical topologies, we refer the reader to the supplementary material. Furthermore, the supplementary material provides face validity assessments demonstrating connection strength inferred by the proposed method correlates significantly over trials with the ground truth.

#### 3.1.1 Network Topology of Neural Interactions Deduced from Hemodynamics

In these results, we performed 100 simulations of the network topology with ground truth illustrated in Fig. 3 and Fig. 4a. In Fig. 4b and c, we see that *ROI*_1_ consistently sends information to *ROI*_2_ and *ROI*_3_ with strong strength of connection. On the other hand, the inferred connection between *ROI*_2_ and *ROI*_3_ is much weaker and not significant according to confidence bounds. However, looking at the contrast network in Fig. 4d, the confidence interval is much smaller for the connection between *ROI*_2_ to *ROI*_3_ and does not include zero. A Wilcoxon signed rank test later revealed that this connection was also the only one in the network with values significantly different between the two conditions.

**Fig 4.**
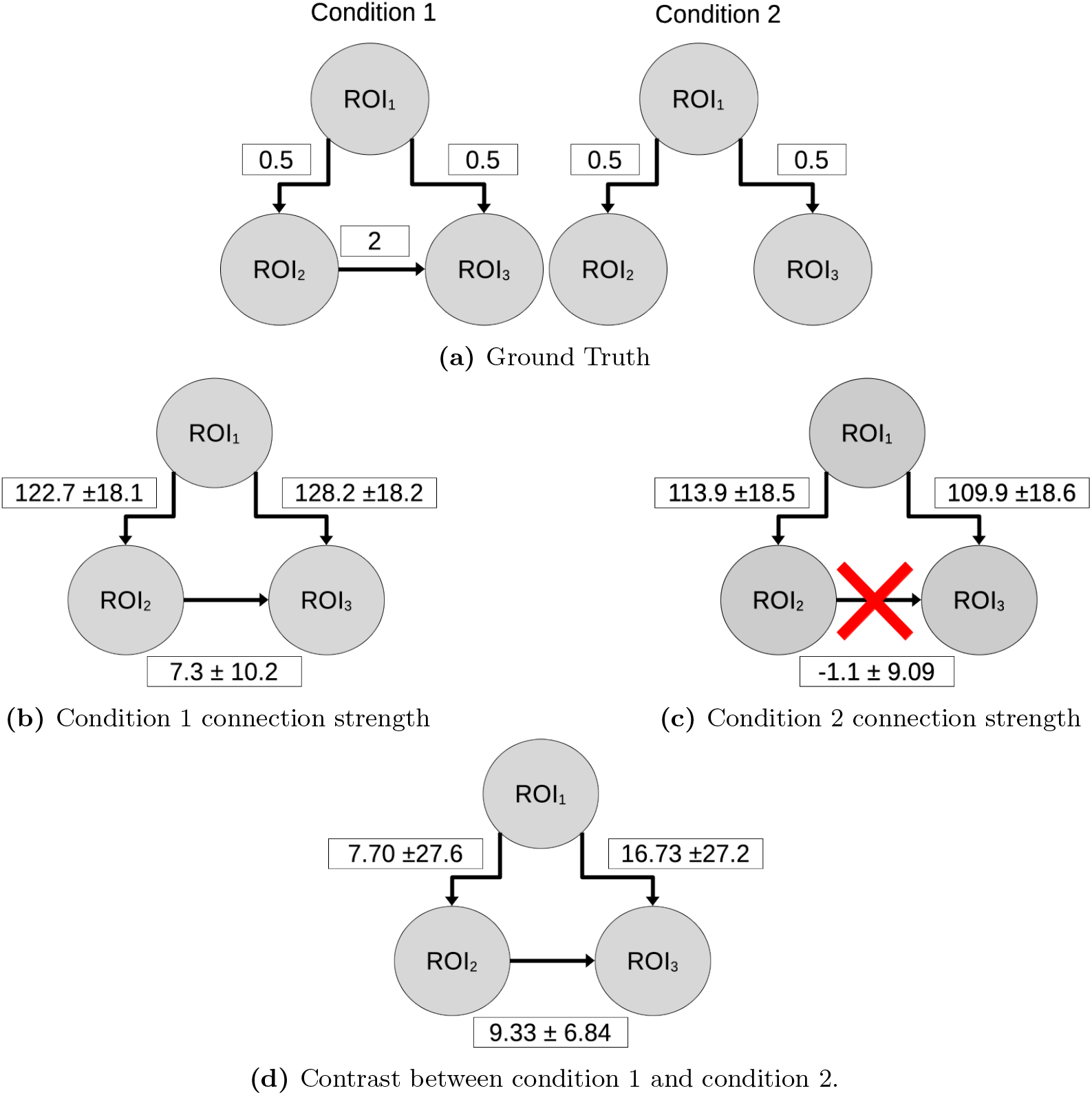
Inferred connection strength from PPI GP-CCM, given ground truth connectivity in (a), are displayed for either condition 1 (b) or condition 2 (c), along with their 95% confidence interval bounds. The contrast between condition 1 and condition 2 can be seen in (d), with 95% confidence interval bounds. Confidence bounds for which zero is not inclusive implies significance. When contrasting between condition 1 and condition 2, where condition 1 has a complete connection from *ROI*_2_ to *ROI*_3_, and condition 1 has a broken connection from *ROI*_2_ to *ROI*_3_, the only connection shown to be significantly different between the two conditions was *ROI*_2_ to *ROI*_3_, as seen in (d).

### 3.2 Construct Validity Tests

After confirming the face validity of the proposed method through simulations, we finally test the construct validity by using a face perception dataset as described on section 2.3.1 to see whether the proposed PPI GP-CCM method is able to uncover DCM effectivity connectivity findings [35].

#### 3.2.1 Face Perception Dataset

To perform task-evoked connectivity analysis, we contrasted the connection strength obtained when participants were presented with faces to the connection strength when participants were presented with arbitrary objects in Fig. 5. In Fig. 5a, we visualize the distribution of contrasting connectivity strength on the group level in comparisons of face stimuli vs shape stimuli. Fig. 5b shows a matrix of T scores for the connectivity analysis on the group level, thresholding results that did not survive multiple comparison correction using Bonferroni. In Fig. 5c, we illustrate nodal interactions in a graphical representation. After multiple comparison corrections, we found significant connectivity in the right hemisphere from the OFA to the FFA and the STS, with weaker, but still significant interactions from the STS to the FFA. These results are in accordance with [35]. The EVC only directed information towards the FFA. In the left hemisphere, we only observed feedforward interactions from the OFA to the FFA and STS, with no intercommunication between the FFA and STS.

**Fig 5.**
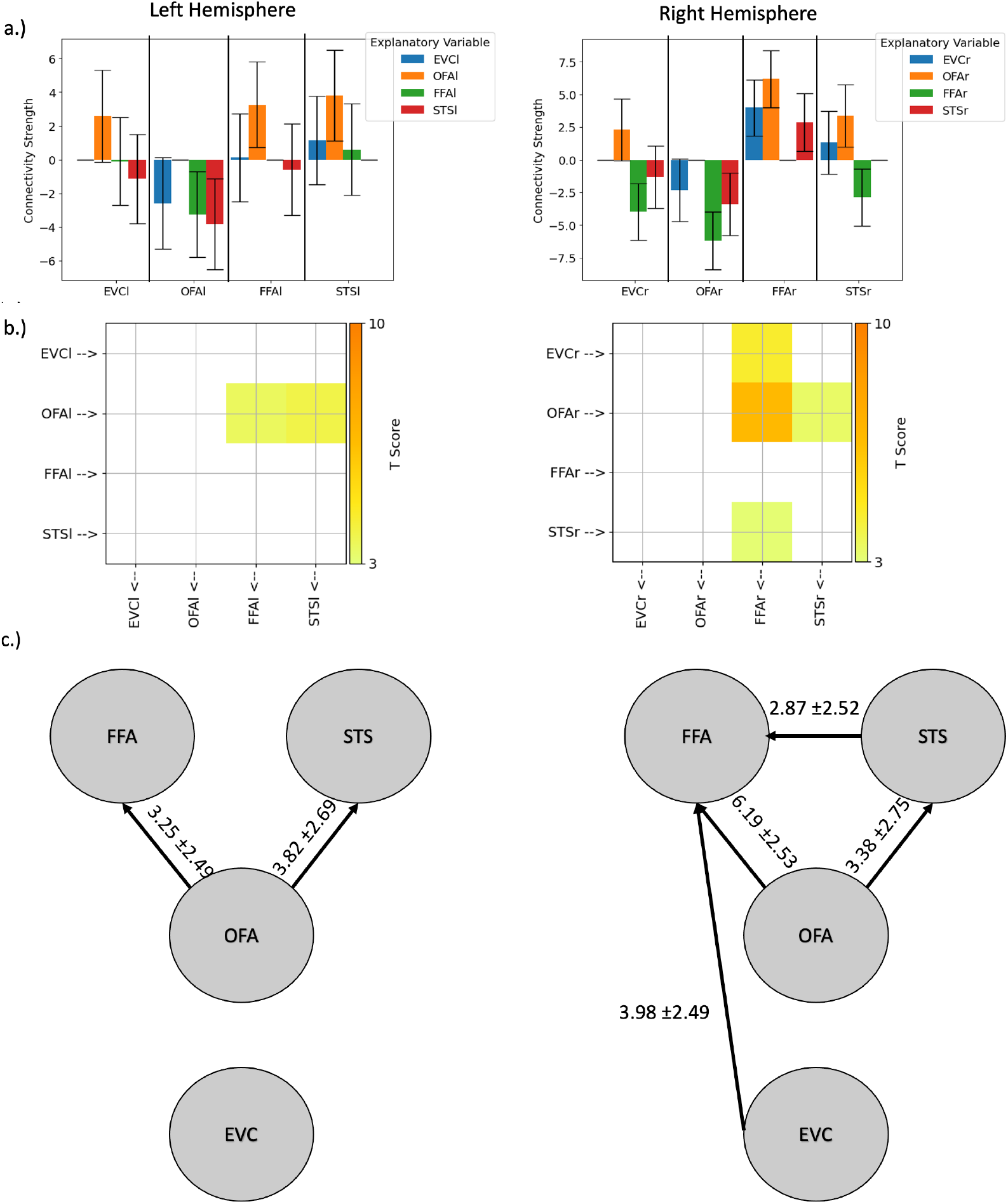
Results of PPI GP-CCM on the HCP emotional faces BOLD dataset, contrasting connectivity during presentation of emotional faces vs. arbitrary shapes at the ROI nodes specified in [35]. a.) Strength of connectivity into different ROIs (x axis) of the left or right hemisphere (separate panels) received from all other ROIs of the same hemisphere (indicated by the colors). Bars are means, errorbars are confidence intervals. b.) T-scores from a group analysis of connections that survived Bonferroni correction for multiple comparisons. c.) Graphical depiction of the nodal interactions between ROIs in the left or right hemisphere (separate panels) that show significantly increased connectivity during presentation of emotional faces compared to shapes. Edges are labeled with the mean connection strength *±* the confidence interval.

## 4 Discussion

In this paper, we describe and test a novel method for uncovering connectivity from BOLD signal data acquired during tasks. Our method takes into consideration contextual dependence in state space by extending our previously described Gaussian Processes Convergent Cross-Mapping method with a nonstationary covariance function for psychophysiological interactions dictated by an (unobserved) physiological signal. We evaluate the strength of evidence for a connection using simulations of neurovascular systems of equations. We then demonstrate the method’s power to perform path analysis: using eq. 7, we assess posterior inference of cross-mapping effects to detect changes in network topology between conditions as illustrated in 3. Lastly, we test whether the proposed method can uncover previously documented effective connectivity during a face perception task.

We evaluated whether our method can infer changes in the acyclic neural population topologies diagrammed in 3 from emergent simulated neurovascular responses. The acyclic graph fulfills the assumptions of the inference of unidirectional flow permitted by the proposed PPI GP-CCM method. We hypothesized that by having one condition with all connections closed and one condition where one connection is open (severed), we should be able to infer which connection was modulated between tasks using statistical contrast methods, i.e. a Wilcoxon signed rank test. Fig. 4d demonstrates that our proposed method succeeded.

This acyclic graph topology presented a trivial case for our proposed method, i.e. statistical inference looking at just the first-level for a single condition would show the effect without any need for contrast statistics. On the other hand, cyclic topologies such as supplementary Fig. 4 are nontrivial as they ostensibly violate a major assumption of unidirectional coupling. As explained on in supplementary material section 3, embeddings of time series preserve only the gradients information needed to generate the time series. In cyclic graphs, all nodes require the full gradient information of the generating system; as each node’s dynamics are dependent on all other variables, cross-mapping would thus never show any information gain as measured by eq. 9. However, if a network topology changes, i.e. from a ring to a chain (by opening up one of the connections), all connections from the node where outgoing connections were disrupted should see a significant change, i.e. outgoing connections are significantly less than zero and incoming connections would be significantly greater than zero. Thus, if an assumption of cyclicity were to be made about network interactions, inference can not be meaningfully made without a contrasting condition. With this in mind, in Supplementary Fig. 5, we present results of contrasting a ring connection before and after a connection was opened, demonstrating the aforementioned effect.

From the pairwise connected populations described in Supplementary Section 1 and Supplementary Table 1, we showed through simulations that Bayes factors exceed substantial strength of evidence. This result indicates that in the event that a positive connectivity is reported, our knowledge about the connectivity increases substantially [52]. While sensitivity analyses yield low values, negative results do not substantially decrease our odds of identifying a connection. For example, the Bayes factor for negative results, i.e. 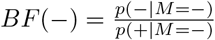 for *ϵ*_*y*_ = 4 : 42%, would be 1.69. This value means that obtaining a negative result would, at most, less than double our confidence about the absence of a connection.

While in all cases, PPI GP-CCM provided substantial evidence for the coupling that we simulated (Supplementary Table 1), we noted that the Bayes Factor was curiously lower in simulations without clear simulated coupling direction (simulation 1 and 3, condition 2) than in simulations with coupling. This may be explained by the sensitivity to difference of entropy during condition 1 may having been reduced significantly when exploring condition 2, however no substantial amount of difference of entropy was observed in condition 2. This may lead the PPI GP-CCM, if sensitive enough, to detect significant changes in a-posteriori entropy leaking from condition 1 to condition 2. This motivates contrast analysis between conditions, with methods such as the Wilcoxon signed rank test that was used to obtain subsequent results on network topologies in simulations and empirical data in this paper.

Using a receiver operating characteristic curve analysis, Supplementary Fig. 2 further demonstrates the power of PPI GP-CCM for detecting true coupling with high sensitivity while maintaining specificity. The result shows that a first-level analysis (at the individual level) may permit higher *α*, substantially increasing sensitivity while keeping subject-level specificity (1 −*FPR*) greater than 50% such that the second level tests for coupling may maintain specificity.

We note that these are simulations where the system’s equations we are comparing can be parcellated exactly, i.e. there are no cross-contaminations between neural populations in the simulations. However, biological systems and fMRI BOLD data contain substantial variance due to mislocalization of neural activity estimated from BOLD signals (BOLD signals are dominated by large blood vessels, which are not located within the active neural tissue) [62]. To avoid deriving overly strong inferences from false positive results, we propose to perform group-level contrast statistics such as a Wilcoxon or T test to assess differences between conditions.

In order to further demonstrate that the method can perform valid contrast statistics, we reutilized the same pairwise interacting population model to establish that the connection strength revealed by PPI GP-CCM follows the trend of ground-truth connection strengths (face validity). As illustrated in Supplementary Fig. 3, we performed simulations with variable connection strengths (5 levels). In simulations 1 and 3 (Supplementary Fig. 3, left panels), event 1 evoked connections from *ROI*_1_ to *ROI*_2_ that increased in strength across levels, while event 2 induces no coupling in the system. The results reflected this: connection strengths assigned to event 2 were consistently distributed around 0, while connection strengths during event 1 increased monotonically with the level of the simulated connection strength. In simulations 2 and 4, event 1 again induced positive connection with a strength that increased over 5 levels. For event 2 however, we simulated the opposite directionality: a negative connection increasing in strength with the connection level. We thus expected more negative estimated connectivity values as connection strength increased for event 2. Again, these patterns were correctly detected by our method (Supplementary Fig. 3, right panels).

In order to assess the construct validity of our method, we aimed to replicate previously documented directed functional connectivity findings about face perception in a well studied dataset, i.e. the WU-Minn Human Connectome S900 Release [28, 35]. Our connectivity analysis, when constrained to the face localized areas in the right hemisphere as determined in [35], revealed that the STS and the FFA received information from OFA, while the EVC and the STS directed information to the FFA, as seen in Fig. 5. This result resembles the feedforward model with greatest log evidence presented in [35], lacking only a connection from the EVC to the STS. However, in 5a, the EVC’s interaction with STS does, in fact, have a positive mean value on the group level. It could be interesting to do further evaluations to determine whether there are subsamples of participants that show or do not show this connection.

Time-lag based methods of connectivity in fMRI analysis have been often discredited considering the sluggish nature of BOLD signals, as well as the nonlinear effects that time-constants may have on delayed neurovascular response [5, 63, 64]. For this reason, generative models in the vein of dynamic causal models have been preferred in neuroimaging literature [42, 43, 65]. GP-CCM and the proposed method in this paper, PPI GP-CCM, attempt to mitigate limitations of time-lag based methods by imposing prior Gaussian process constraints to lift time-lag methods to a generative statistical model of the state space manifold for nonparametric model evidence comparisons. As formulated here, this results in combinatorial pair-wise model evidence comparisons between regions of interests, i.e. parcellations of the cortex in our case. However, it is understood that the brain contains a slew of recurrent networks [47–49], paths of information that result in indirect coupling [50], and phenomena emerging from high-order statistical behaviors such as redundancy and synergy [66, 67]. We indeed take mediating interactions into consideration in the model, however to further understand complex networks, higher order interactions, such as those on hypergraphs, will need to be integrated into the proposed method. On that note, future work to overcome these limitations may be directed towards looking at spatial scale of interactions on networks, for example by performing path analysis based on a combination of interactions between multiple nodes, to evaluate if that combination better explains interaction with another set of nodes.

## 5 Conclusion

We introduce and validate a novel method that augments the covariance function in GP-CCM to enable task-dependent connectivity analysis from BOLD data, leveraging psychophysiological interaction theory. Our method demonstrates substantial strength of evidence regarding directional connectivity between pairs of regions, as demonstrated through simulations of coupled chaotic systems and through neural networks observed by hemodynamics. Additionally, we demonstrate the utility of our proposed method by uncovering previously described connections associated with face perception in the ventrolateral occipito-temporal cortex.

Our proposed method represents a promising tool for performing pairwise directional influence tests to uncover interactions between regions of interest in the brain. Overall, our study highlights the potential of our method to improve our understanding of functional connectivity between brain regions, particularly in the context of task-dependent neural responses. Future research may seek to apply our method to explore additional brain networks and cognitive processes, building on our results to further advance our understanding of the human brain.

## Supporting information

Supplemental material/results

## Acknowledgements

This study was financed in part by the European Commission Horizon 2020 project RHUMBO, grant agreement ID 813234, DOI: 10.3030/813234, and by the Italian Ministry of Education and Research (MIUR) in the framework of the ForeLab project (Departments of Excellence).

## Data and Code Availability Statement

All code used for simulating, obtaining, or processing data can be found in the following GitHub repository: https://github.com/lemiceterieux/PPI-GP-CrossMapping.

## References

1. Gatica M, Cofré R, Mediano PAM, Rosas FE, Orio P, Diez I, et al. High-Order Interdependencies in the Aging Brain. Brain Connectivity. 2021;11(9):734–744.

2. Friston KJ, Price CJ. Degeneracy and redundancy in cognitive anatomy. Trends in Cognitive Sciences. 2003;7(4):151–152.

3. Latham PE, Nirenberg S. Synergy, Redundancy, and Independence in Population Codes, Revisited. Journal of Neuroscience. 2005;25(21):5195–5206.

4. Stramaglia S, Angelini L, Wu G, Cortes JM, Faes L, Marinazzo D. Synergetic and Redundant Information Flow Detected by Unnormalized Granger Causality: Application to Resting State fMRI. IEEE Transactions on Biomedical Engineering. 2016;63(12):2518–2524. doi:10.1109/TBME.2016.2559578.

5. Friston KJ. Functional and effective connectivity: a review. Brain connectivity. 2011;1(1):13–36. doi:10.1089/brain.2011.0008.

6. Friston KJ. Functional and effective connectivity in neuroimaging: a synthesis. Human brain mapping. 1994;2(1-2):56–78.

7. Leonardi N, Van De Ville D. On spurious and real fluctuations of dynamic functional connectivity during rest. Neuroimage. 2015;104:430–436.

8. Zalesky A, Breakspear M. Towards a statistical test for functional connectivity dynamics. Neuroimage. 2015;114:466–470.

9. Marrelec G, Krainik A, Duffau H, Pélégrini-Issac M, Lehéricy S, Doyon J, et al. Partial correlation for functional brain interactivity investigation in functional MRI. Neuroimage. 2006;32(1):228–237.

10. Friston KJ, Buechel C, Fink GR, Morris J, Rolls E, Dolan RJ. Psychophysiological and Modulatory Interactions in Neuroimaging. NeuroImage. 1997;6(3):218–229.

11. O’Reilly JX, Woolrich MW, Behrens TE, Smith SM, Johansen-Berg H. Tools of the trade: psychophysiological interactions and functional connectivity. Social cognitive and affective neuroscience. 2012;7(5):604–609.

12. Gitelman DR, Penny WD, Ashburner J, Friston KJ. Modeling regional and psychophysiologic interactions in fMRI: the importance of hemodynamic deconvolution. Neuroimage. 2003;19(1):200–207.

13. Friston K. Dynamic causal modeling and Granger causality Comments on: the identification of interacting networks in the brain using fMRI: model selection, causality and deconvolution. NeuroImage. 2011;58(2):303–311.

14. Frässle S, Lomakina EI, Razi A, Friston KJ, Buhmann JM, Stephan KE. Regression DCM for fMRI. Neuroimage. 2017;155:406–421.

15. Zhou Z, Wang X, Klahr NJ, Liu W, Arias D, Liu H, et al. A conditional Granger causality model approach for group analysis in functional magnetic resonance imaging. Magnetic resonance imaging. 2011;29(3):418–433.

16. Wen X, Yao L, Liu Y, Ding M. Causal interactions in attention networks predict behavioral performance. Journal of Neuroscience. 2012;32(4):1284–1292.

17. Ding M, Bressler SL, Yang W, Liang H. Short-window spectral analysis of cortical event-related potentials by adaptive multivariate autoregressive modeling: data preprocessing, model validation, and variability assessment. Biol Cybern. 2000;83(1):35–45.

18. Friston K. Causal modelling and brain connectivity in functional magnetic resonance imaging. PLoS biology. 2009;7(2):e1000033.

19. Smirnov D, Bezruchko B. Spurious causalities due to low temporal resolution: Towards detection of bidirectional coupling from time series. EPL (Europhysics Letters). 2012;100(1):10005.

20. Smirnov DA. Spurious causalities with transfer entropy. Phys Rev E. 2013;87:042917. doi:10.1103/PhysRevE.87.042917.

21. Sauer T, Yorke JA, Casdagli M. Embedology. Journal of Statistical Physics. 1991;65(3):579–616. doi:10.1007/BF01053745.

22. Takens F. Detecting strange attractors in turbulence. In: Rand D, Young LS, editors. Dynamical Systems and Turbulence, Warwick 1980. Berlin, Heidelberg: Springer Berlin Heidelberg; 1981. p. 366–381.

23. Rasmussen CE, Williams CKI. Gaussian Processes for Machine Learning (Adaptive Computation and Machine Learning). The MIT Press; 2005.

24. Sugihara G, May R, Ye H, Hsieh Ch, Deyle E, Fogarty M, et al. Detecting Causality in Complex Ecosystems. Science. 2012;338(6106):496–500. doi:10.1126/science.1227079.

25. Ghouse A, Faes L, Valenza G. Inferring directionality of coupled dynamical systems using Gaussian process priors: Application on neurovascular systems. Phys Rev E. 2021;104:064208. doi:10.1103/PhysRevE.104.064208.

26. Ghouse A, Valenza G. Inferring parsimonious coupling statistics in nonlinear dynamics with variational gaussian processes. In: Complex Networks and Their Applications XI: Proceedings of The Eleventh International Conference on Complex Networks and Their Applications: COMPLEX NETWORKS 2022— Volume 1. Springer; 2023. p. 377–389.

27. Ahmed, N.A. and Gokhale, D.V. Entropy expressions and their estimators for multivariate distributions IEEE Transactions on Information Theory.1989; 35(3):688–692

28. Van Essen DC, Smith SM, Barch DM, Behrens TE, Yacoub E, Ugurbil K, et al. The WU-Minn human connectome project: an overview. Neuroimage. 2013;80:62–79.

29. Fairhall SL, Ishai A. Effective connectivity within the distributed cortical network for face perception. Cerebral cortex. 2007;17(10):2400–2406.

30. Haxby JV, Hoffman EA, Gobbini MI. The distributed human neural system for face perception. Trends in cognitive sciences. 2000;4(6):223–233.

31. Haxby JV, Gobbini MI, Furey ML, Ishai A, Schouten JL, Pietrini P. Distributed and overlapping representations of faces and objects in ventral temporal cortex. Science. 2001;293(5539):2425–2430.

32. Gschwind M, Pourtois G, Schwartz S, Van De Ville D, Vuilleumier P. White-matter connectivity between face-responsive regions in the human brain. Cerebral cortex. 2012;22(7):1564–1576.

33. Pyles JA, Verstynen TD, Schneider W, Tarr MJ. Explicating the face perception network with white matter connectivity. PloS one. 2013;8(4):e61611.

34. Davies-Thompson J, Andrews TJ. Intra-and interhemispheric connectivity between face-selective regions in the human brain. Journal of neurophysiology. 2012;108(11):3087–3095.

35. Wang Y, Metoki A, Smith DV, Medaglia JD, Zang Y, Benear S, et al. Multimodal mapping of the face connectome. Nature human behaviour. 2020;4(4):397–411.

36. Kessler R, Rusch KM, Wende KC, Schuster V, Jansen A. Revisiting the effective connectivity within the distributed cortical network for face perception. NeuroImage: Reports. 2021;1(4):100045.

37. MacKay DJC. In: Domany E, van Hemmen JL, Schulten K, editors. Bayesian Methods for Backpropagation Networks. New York, NY: Springer New York; 1996. p. 211–254. Available from: https://doi.org/10.1007/978-1-4612-0723-8_6.

38. Friston KJ, Jezzard P, Turner R. Analysis of functional MRI time-series. Human brain mapping. 1994;1(2):153–171.

39. Deshpande G, LaConte S, James GA, Peltier S, Hu X. Multivariate Granger causality analysis of fMRI data. Human brain mapping. 2009;30(4):1361–1373.

40. Seth AK, Barrett AB, Barnett L. Granger causality analysis in neuroscience and neuroimaging. Journal of Neuroscience. 2015;35(8):3293–3297.

41. Roebroeck A, Formisano E, Goebel R. The identification of interacting networks in the brain using fMRI: model selection, causality and deconvolution. Neuroimage. 2011;58(2):296–302.

42. Friston KJ, Harrison L, Penny W. Dynamic causal modelling. NeuroImage. 2003;19(4):1273 –1302. doi:https://doi.org/10.1016/S1053-8119(03)00202-7.

43. Stephan KE, Weiskopf N, Drysdale PM, Robinson PA, Friston KJ. Comparing hemodynamic models with DCM. Neuroimage. 2007;38(3):387–401.

44. Snelson E, Ghahramani Z. Sparse Gaussian Processes Using Pseudo-Inputs. In: Proceedings of the 18th International Conference on Neural Information Processing Systems. NIPS’05. Cambridge, MA, USA: MIT Press; 2005. p. 1257–1264.

45. Aggarwal CC, Hinneburg A, Keim DA. On the Surprising Behavior of Distance Metrics in High Dimensional Space. In: Van den Bussche J, Vianu V, editors. Database Theory — ICDT 2001. Berlin, Heidelberg: Springer Berlin Heidelberg; 2001. p. 420–434.

46. Nevo B. Face validity revisited. Journal of Educational Measurement. 1985;22(4):287–293.

47. Garrido MI, Kilner JM, Kiebel SJ, Friston KJ. Evoked brain responses are generated by feedback loops. Proceedings of the National Academy of Sciences. 2007;104(52):20961–20966.

48. Dehaene S, Changeux JP. Experimental and theoretical approaches to conscious processing. Neuron. 2011;70(2):200–227.

49. Tononi G, Edelman GM. Consciousness and complexity. science. 1998;282(5395):1846–1851.

50. Ross LN. Causal concepts in biology: How pathways differ from mechanisms and why it matters. The British Journal for the Philosophy of Science. 2020;.

51. Glover GH. Deconvolution of Impulse Response in Event-Related BOLD fMRI1. NeuroImage. 1999;9(4):416–429.

52. Jeffreys H. The theory of probability. OUP Oxford; 1998.

53. Kass RE, Raftery AE. Bayes Factors. Journal of the American Statistical Association. 1995;90(430):773–795. doi:10.2307/2291091.

54. Duchaine B, Yovel G. A revised neural framework for face processing. Annual review of vision science. 2015;1:393–416.

55. Sorger B, Goebel R, Schiltz C, Rossion B. Understanding the functional neuroanatomy of acquired prosopagnosia. Neuroimage. 2007;35(2):836–852.

56. Logothetis NK. What we can do and what we cannot do with fMRI. Nature. 2008;453(7197):869–878.

57. Cronbach LJ, Meehl PE. Construct validity in psychological tests. Psychological bulletin. 1955;52(4):281.

58. Hariri AR, Tessitore A, Mattay VS, Fera F, Weinberger DR. The amygdala response to emotional stimuli: a comparison of faces and scenes. Neuroimage. 2002;17(1):317–323.

59. Barch DM, Burgess GC, Harms MP, Petersen SE, Schlaggar BL, Corbetta M, et al. Function in the human connectome: task-fMRI and individual differences in behavior. Neuroimage. 2013;80:169–189.

60. Nguyen VT, Breakspear M, Cunnington R. Fusing concurrent EEG–fMRI with dynamic causal modeling: Application to effective connectivity during face perception. Neuroimage. 2014;102:60–70.

61. McLaren DG, Ries ML, Xu G, Johnson SC. A generalized form of context-dependent psychophysiological interactions (gPPI): a comparison to standard approaches. Neuroimage. 2012;61(4):1277–1286.

62. Hillman EM. Coupling mechanism and significance of the BOLD signal: a status report. Annual review of neuroscience. 2014;37:161.

63. Friston KJ, Bastos AM, Oswal A, van Wijk B, Richter C, Litvak V. Granger causality revisited. NeuroImage. 2014;101:796–808. doi:10.1016/j.neuroimage.2014.06.062.

64. Deshpande G, Sathian K, Hu X. Effect of hemodynamic variability on Granger causality analysis of fMRI. Neuroimage. 2010;52(3):884–896.

65. Frässle S, Harrison SJ, Heinzle J, Clementz BA, Tamminga CA, Sweeney JA, et al. Regression dynamic causal modeling for resting-state fMRI. Human brain mapping. 2021;42(7):2159–2180.

66. Battiston F, Amico E, Barrat A, Bianconi G, Ferraz de Arruda G, Franceschiello B, et al. The physics of higher-order interactions in complex systems. Nature Physics. 2021;17(10):1093–1098.

67. Luppi AI, Mediano PA, Rosas FE, Holland N, Fryer TD, O’Brien JT, et al. A synergistic core for human brain evolution and cognition. Nature Neuroscience. 2022;25(6):771–782.

