## Supplemental material/results for "Estimation of Task-Evoked Directed Functional Connectivity by Cross-Mapping Psychophysiological Variables"

#### 1 Simulations of Pairwise-coupled Neurovascular Neural Populations

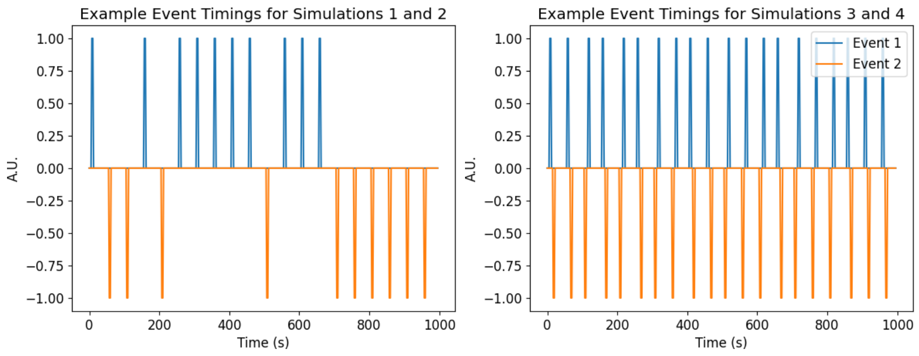


(a)

Figure 1: Event timings for simulations 1 and 2 (a) and simulations 3 and 4 (b). in the pairwise-coupled neurovascular neural populations

In order to assess the validity of the proposed PPI GP-CCM on metabolic signals related to neural activity as obtained using functional Near InfraRed Spectroscopy (fNIRS) or functional Magnetic Resonance Imaging (fMRI), we exploit forward models of hemodynamics to simulate neurovascular signals emerging from neural networks [43]. In these simulations, we have populations 1 and 2, each comprising 50 neural units. As per [43], a bilinear state equation is used to model neural activity signals ($x$) induced by a cognitive task:

$$\dot{x} = Ax + \sum_{j=1}^{m} u_j B^j x + Cu \qquad (1)$$

The $A$ matrix describes the autonomous dynamics of the system; in our simulations, we treat it as a diagonal matrix with values -1, implying dynamics returning to a baseline 0 state. $u$ are events of a cognitive condition (where the element $u_j$ is 0 when $jth$ condition is not occurring, and 1 when the $j^{th}$ condition is occurring). $B^j$ describes the interconnections between populations of interest (i.e. $\begin{bmatrix} R_{11} & R_{12} \\ 0 & R_{22} \end{bmatrix}$ for population 1 coupled towards population 2, and each $R_{ij}$ is an i.i.d. random matrix $R_{ij} \sim \mathcal{MN}(0, 0.4, 1)$). The B matrices do not change in time, rather they are realized for each simulated time.

We perform 4 different simulations with three conditions $u_1$, $u_2$ and $u_3$. For simulation 1, $u_1$ and $u_2$ are events that occur 10 times each over a period of 1000 seconds; each event lasts 6 seconds, events occur at intervals of 1 minute. $u_1$ invokes population 1 to population 2 interactions via a random matrix, $R^1_{12} \sim \mathcal{MN}(0, 0.4, 1)$ while $u_2$ does not invoke interaction between the populations, i.e. $R^2_{12} = 0$. For simulation 2, $R^2_{12} \sim \mathcal{MN}(0, 0.4, 1)$ is instead inducing connections from population 2 to population 1. Event timings can be seen in fig. 1a.

In simulation 3 and 4, $u_1$ and $u_2$ are events that occur 20 times with stimulus duration of 6 seconds, and a short interevent time interval of 4 seconds (within the traditional peak time of the canonical linear hemodynamic response function [51]). Each pair of events are separated by 1 minute. The order of events are randomized for each pairing. Similar to simulations 1 and 2, simulations 3 and 4 either have $u_2$ not induce any interactions between populations, or have $u_2$ inducing connectivity from population 2 to population 1 respectively. Event timings can be seen in fig. 2Event timings for the conditions being simulatedfigure.caption.4b. For all simulations, we also aimed to evaluate the effect of true connection strength on the statistic of coupling strength, and whether it indeed correlates and follows expected trends. We multiply the antidiagonal elements of the connectivity matrices by the desired connectivity, with connectivity values {0,0.2,0.4,0.8}.

We perform 4 different simulations with three conditions $u_1$, $u_2$ and $u_3$. For simulation 1, $u_1$ and $u_2$ are events that occur 10 times each over a period of 1000 seconds; each event lasts 6 seconds, events occur at intervals of 1 minute. $u_1$ invokes population 1 to population 2 interactions via a random matrix, $R^1_{12} \sim \mathcal{MN}(0, 0.4, 1)$ while $u_2$ does not invoke interaction between the populations, i.e. $R^2_{12} = 0$. For simulation 2, $R^2_{12} \sim \mathcal{MN}(0, 0.4, 1)$ is instead inducing connections from population 2 to population 1. Event timings can be seen in fig. 2Event timings for the conditions being simulatedfigure.caption.4a.

In simulation 3 and 4, $u_1$ and $u_2$ are events that occur 20 times with stimulus duration of 6 seconds, and a short interevent time interval of 4 seconds (within the traditional peak time of the canonical linear hemodynamic response function [51]). Each pair of events are separated by 1 minute. The order of events are randomized for each pairing. Similar to simulations 1 and 2, simulations 3 and 4 either have $u_2$ not induce any interactions between populations, or have $u_2$ inducing connectivity from population 2 to population 1 respectively. Event timings can be seen in fig. 2Event timings for the conditions being simulatedfigure.caption.4b.

### 2 Pairwise Coupling of Neural Populations Deduced from Hemodynamic Responses

| | Condition 1 | | | Condition 2 | | | |
|---|---|---|---|---|---|---|---|
| $N_{simuls} = 100$ | $Pop_1 \to Pop_2$ | $Pop_2 \to Pop_1$ | $BF_{12}$ | $Pop_1 \to Pop_2$ | $Pop_2 \to Pop_1$ | $BF_{21}$ | $BF_{any}$ |
| Sim 1 | 77 | 1 | 77.0 | 35 | 17 | 0.48 | 4.35 |
| Sim 2 | 76 | 4 | 19.0 | 1 | 78 | 78.0 | 30.8 |
| Sim 3 | 74 | 4 | 18.5 | 39 | 19 | 0.48 | 3.58 |
| Sim 4 | 75 | 1 | 75.0 | 3 | 78 | 26.0 | 38.3 |

Table 1: Results of face validity tests aiming to detect the coupling direction of pairwise coupled neural populations from hemodynamics. Values in the table are N reported connections for each direction out of 100 realizations of a given simulation. In Condition 1, all simulations contained induced coupling from network 1 ($Pop_1$) to network 2 ($Pop_2$). In Condition 2, Sim 1 and Sim 3 contain no coupling, while Sim 2 and 4 contain coupling from $Pop_2$ to $Pop_1$. $BF_{any}$ is calculated from the ratio of correct inferred direction to incorrectly inferred direction.

The results of the simulations (Table 1) show that the number of times our method detects the simulated connection direction is much higher than the number of times if reports the opposite connection direction (see fig. 2Event timings for the conditions being simulatedfigure.caption.4 for the simulated event time courses). For example, in Condition 1 where there always was a connection from $Pop_1$ to $Pop_2$, connections with this direction are reported much more often than connections in the opposite direction, yielding high Bayes factors ($BF_1 2$). Considering there were 100 realizations for each simulation, the sensitivity appears generally over 70%. The worst specificity achieved for any of the simulations was 79%. As evidenced by the Bayes factors, according to standard terminology [52], a positive value from our model is strong evidence ($BF > 10$) for a connection. Simulated directed connections between two neural populations can be detected irrespective of the connection direction (see Sim 2 and 4). We note however that when an event induces no directional connection between the networks (Condition 2 in Sim 1 and 3), the Bayes factor still indicates substantial evidence according to [52].

In fig. 2 we demonstrate the sensitivity curve as a function of the first level significance level parameter $\alpha$. In all simulations, we demonstrate an area under the curve (AUC) of over 0.7 for sensitivity of true coupling direction, while the false coupling detection all have an AUC of less than 0.5.

Finally, in fig. 3, we demonstrate the trend of the proposed method, PPI

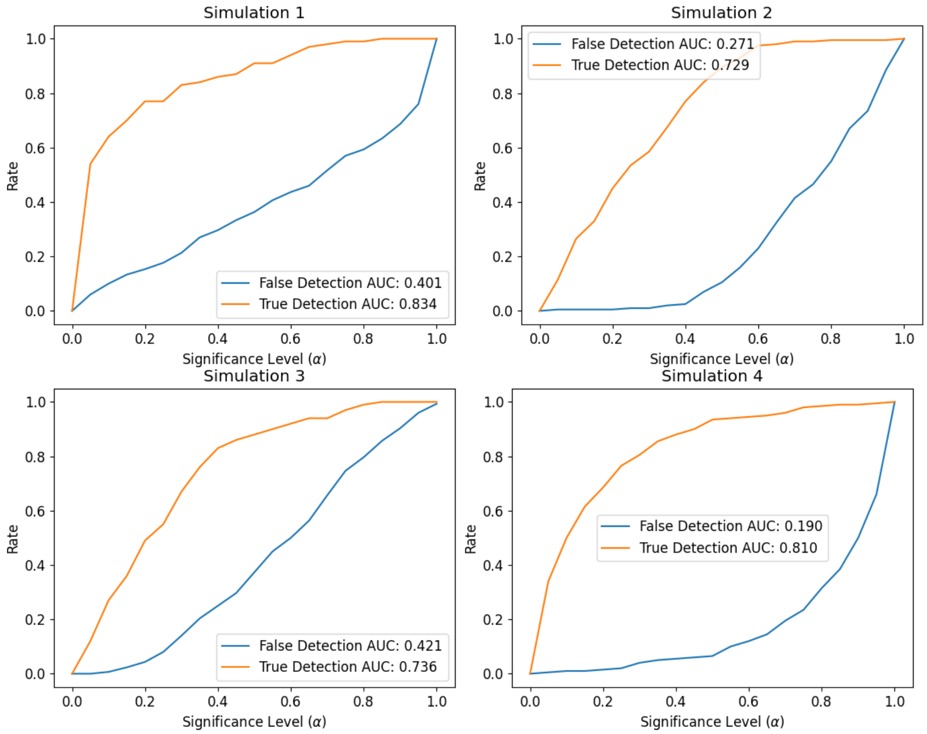


Figure 2: Sensitivity curves as a function of the first level significance level $\alpha$ for simulations 1 (a), 2 (b), 3 (c) and 4 (d). On the X-axis, the significance threshold of the permutation test is varied from 0 to 1, and on the Y axis, the rate at which either a true coupling detection (orange) or a false coupling detection (blue) are reported.

GP-CCM, with respect to increasing connectivity strength. Negative values correspond to an $ROI_2$ forcing $ROI_1$, while a positive value implies the reverse direction. We see that, regardless of interevent interval duration, the method follows expected increase of inferred connection strength for simulation 2 and 4. For simulations 1 and 3, we would expect event 2 to cause no coupling in either direction according to the simulations, which is further reflected in the inferred connection strength discovered by PPI GP-CCM.

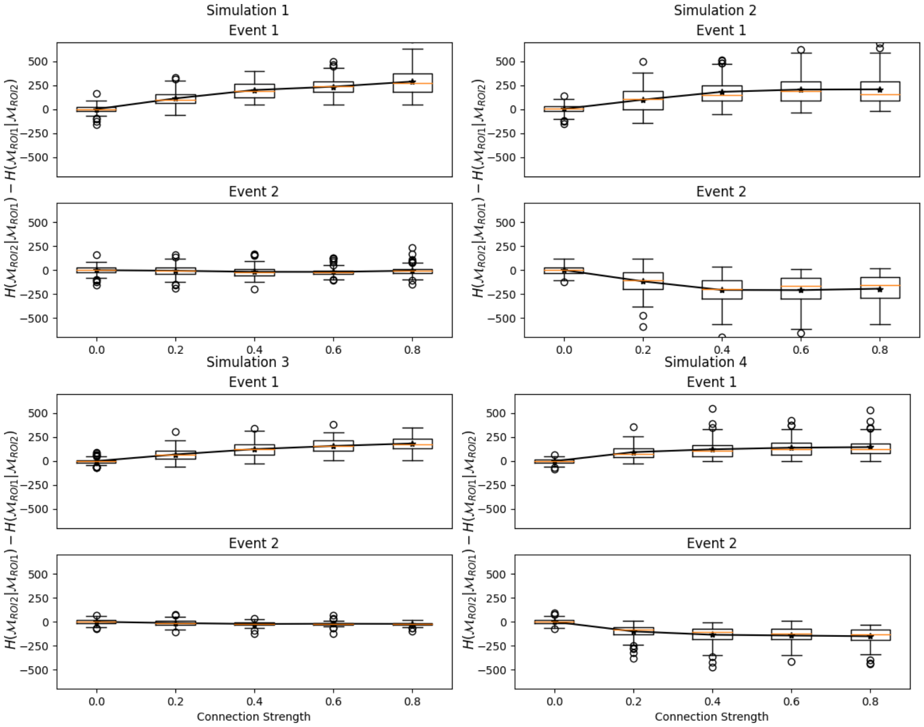


Figure 3: Inferred connection strength statistic compared to the true connection strength for simulations 1 (a), 2 (b), 3 (c) and 4 (d). On the X-axis, the connection strength is varied according to the model in eq. 1, and on the Y axis the connection strength reported by the connectivity statistic. Using the logic that the embedding of a forced variable should be innovating upon states of an embedding of a putative variable, if $H(\mathcal{M}_{ROI2}|\mathcal{M}_{ROI1}) > H(\mathcal{M}_{ROI1}|\mathcal{M}_{ROI2})$, $ROI_1$ would be inferred to be coupled towards $ROI_2$. In event 1, in all simulations, $ROI_1$ should be coupled towards $ROI_2$, thus the statistic should present a positive valued result. In event two, only for simulations 2 and 4, do we induce coupling from $ROI_2$ to $ROI_1$, thus in the inferred connection strength should be negative valued. Otherwise, the result should be zero valued, indicating no significant inferred coupling.

### 3 Cyclic Toplogies

We also examine cyclic ring topologies fit with recurrent interactions (see fig. 4). In other words, the topology includes feedback where all nodes inherently force all other nodes, while at the same time the nodes self-interact indirectly. This topology would, without any proper statistical contrast, pose great difficulty for making inferences with the proposed method. Cross-mapping measures hinge upon embedding theorems that guarantee a preservation of necessary gradient information of the original system that generates the observed time series [21, 22, 24].

If all variables are connected in a cyclic ring formation where a loop of interactions can be made from a node back to itself (condition 1), that would mean no coupling inference can be made as all reconstructed state spaces using an embedding should have the same information, even when taking mediators into consideration. In other words, cross-mapping would provide no surprise. However, if this ring were to be opened (condition 2), e.g. from the dashed connection $ROI_2$ to $ROI_3$ as illustrated in the fig. 4, we hypothesized that we should be able to see significantly less information directed from $ROI_2$ to all other nodes as the influence of $ROI_2$ on those nodes is cut from the graph (the ring becomes a chain terminating at $ROI_3$). This alternatively implies that all information being directed into $ROI_2$ from all other nodes should be significantly increased. Thus, with this simulation, we highlight that although the mathematical assumptions of the Bayesian inference model will be violated in this topology, properly formed statistical tests for contrast should still provide meaningful results about the functional interactions between nodes in the graph. We emphasize the point "functional", as the effect of opening the connection on far nodes has a cascade effect without being direct. This would be similar, as an illustrative example, as stating that the fusiform gyrus functionally couples with the superior temporal sulcus to parse variable face features (such as facial expression) without direct structural connection, but instead through mediated interactions in the occipital face area (inferior occipital gyrus) or other proposed streams [54, 55]. Note that we use this example without claiming that this is a true state of affairs.

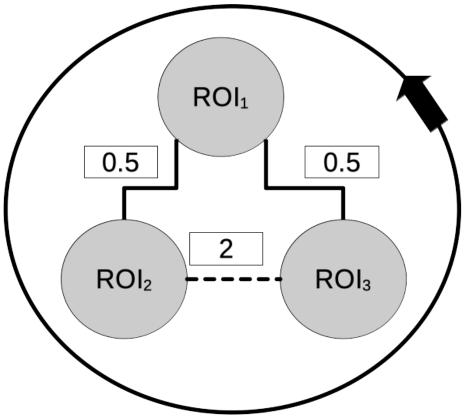


Figure 4: Cyclic functional interactions: In condition 1 the ring is closed such that all ROIs directly or indirectly have a path of interaction with all ROIs including themselves, whereas in condition 2 the ring is open, turning the topology into a chain of interactions that begins at $ROI_3$ and terminates at $ROI_2$. The connection strength parameters for the simulations are written above the respective connections

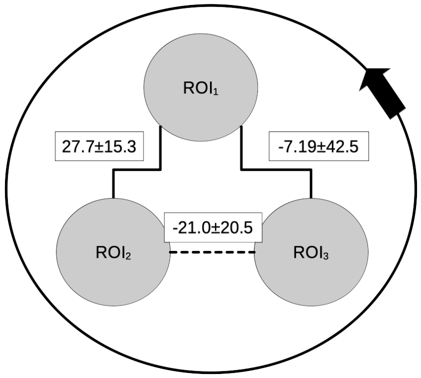


Figure 5: Contrast network between condition 1 (closed connection at dashed line) and condition 2 (opened connection at dashed line) for the ring topology. Reported numbers are the mean alongside the 95% confidence interval bounds. Only connections interacting with $ROI_2$ were found to be significantly different from zero when contrasting condition 1 and condition 2, where outgoing connections were significantly less than zero when opening the connection, while incoming connections were significantly greater than zero. We only show results of contrast to avoid misinterpretation of results on ring topologies.

We then simulated the topology seen in fig. 4, where fig. 5 displays the contrast between condition 1 (closed ring) and condition 2 (open ring). It can be seen that only at connections coming into and out of $ROI_2$ are significantly different from zero, as confirmed by both the 95% confidence interval and a Wilcoxon signed rank test. Connections coming into $ROI_2$ are significantly greater, while connections coming out of $ROI_2$ are significantly less, following our proposed hypothesis regarding information interruption in cyclic networks gleaned from cross-mapping condition contrasts. In fig. 5, we only demonstrate the contrast between network connections rather than connection strength for each condition to avoid misinterpretation of the results, as inference directly on the ring condition can not be made without contrasts given the recurrent connections.
